# Engineered variants provide new insight into the structural properties important for activity of the highly dynamic, trimeric protein disulfide isomerase, PmScsC

**DOI:** 10.1101/421420

**Authors:** Emily J. Furlong, Fabian Kurth, Lakshmanane Premkumar, Andrew E. Whitten, Jennifer L. Martin

**Affiliations:** Institute for Molecular Bioscience, The University of Queensland, St Lucia, QLD 4072, Australia.; Griffith Institute for Drug Discovery, Griffith University, Nathan, QLD 4111 Australia.; Australian Nuclear Science and Technology Organisation, Lucas Heights, NSW 2234, Australia.

## Abstract

Suppressor of copper sensitivity protein C from *Proteus mirabilis* (PmScsC) is a homotrimeric disulfide isomerase that plays a role in copper tolerance – a key virulence trait of the uropathogen. Each protomer of the enzyme has an N-terminal trimerisation stem (59 residues) containing a flexible linker (11 residues) connected to a thioredoxin-fold-containing catalytic domain (163 residues). Here, we characterise two PmScsC variants, PmScsCΔN and PmScsCΔLinker. PmScsCΔN, is an N-terminally truncated form of the protomer with two helices of the trimerisation stem removed, generating a protein with dithiol oxidase rather than disulfide isomerase activity. The crystal structure of PmScsCΔN reported here reveals – as expected – a monomer that is structurally similar to the catalytic domain of native PmScsC. The second variant PmScsCΔLinker was designed to remove the 11 amino acid linker and we show that it generates a protein that has neither disulfide isomerase nor dithiol oxidase activity. The crystal structure of PmScsCΔLinker reveals a trimeric arrangement, with the catalytic domains packed together very closely. Small angle X-ray scattering analysis found that native PmScsC is predominantly trimeric in solution even at low concentration, whereas PmScsCΔLinker exists as an equilibrium between monomeric, dimeric and trimeric states, with the monomeric form dominating at low concentrations. These findings increase our understanding of disulfide isomerase activity, showing how (i) oligomerisation, (ii) spacing between, and (iii) dynamic motion of, catalytic domains in PmScsC all contribute to its native function.

## 1. Introduction

Protein disulfide isomerases are enzymes that proofread and shuffle incorrect disulfide bonds in misfolded protein substrates, and are important for the correct folding and function of many secreted proteins (Berkmen *et al.*, 2005, Hiniker & Bardwell, 2004). The prototypical bacterial disulfide isomerase is Disulfide bond (Dsb) protein C from *Escherichia coli* (EcDsbC) (Zapun *et al.*, 1995, Missiakas *et al.*, 1994, Shevchik *et al.*, 1994). EcDsbC functions as a dimer; when the N-terminal dimerisation domain is deleted the resulting protein lacks disulfide isomerase activity (Sun & Wang, 2000) and it is also unable to interact with its redox partner EcDsbD (Goldstone *et al.*, 2001). Each EcDsbC protomer has a thioredoxin-fold catalytic domain with a redox active motif consisting of two cysteines separated by two other amino acids (CXXC) (McCarthy *et al.*, 2000). The disulfide isomerase activity of EcDsbC requires that the catalytic cysteines are in the dithiol reduced form (Darby *et al.*, 1998) and this form is generated by interaction of EcDsbC with its redox partner membrane protein EcDsbD (Goldstone *et al.*, 2001).

Recently, we reported the structural and functional characterisation of *Proteus mirabilis* suppressor of copper sensitivity protein C (PmScsC) (Furlong *et al.*, 2017). Like DsbC, PmScsC is a protein disulfide isomerase with a thioredoxin-fold catalytic domain (Furlong *et al.*, 2017), redox-active cysteines (Furlong *et al.*, 2017), and a redox partner membrane protein (PmScsB) that reduces the active site cysteines (Furlong *et al.*, 2018). Unlike DsbC, PmScsC is trimeric rather than dimeric (Furlong *et al.*, 2017). Three crystal structures and SAXS analysis also revealed that PmScsC is highly dynamic (Furlong *et al.*, 2017). The region responsible for this conformational flexibility is an 11 amino acid linker within the trimerisation stem of the protein. Replacement of the flexible linker with a rigid helical peptide linker (PmScsC RHP), limits the rigid body motion of the catalytic domain relative to the trimer stalk, and generates a protein that is inactive as a disulfide isomerase. Moreover, PmScsC lacking the trimerisation stem (N-terminal 41 residues, PmScsCΔN) is also inactive as a disulfide isomerase but does have dithiol oxidase activity (Furlong *et al.*, 2017). Neither the native protein, nor the PmScsC RHP variant, exhibited dithiol oxidase activity. The difference in activity profile between the native protein (disulfide isomerase) and PmScsCΔN (dithiol oxidase) was proposed to be a consequence of their different oligomeric states.

Here we report the structural characterisation of PmScsCΔN and the structural and functional characterisation of PmScsCΔLinker (a variant in which the 11-amino acid flexible linker is deleted, Figure 1). The crystal structure of PmScsCΔN reveals – as expected – a monomeric protein that closely resembles the catalytic domain from the full-length PmScsC protomer. We show that the PmScsCΔLinker variant lacks both disulfide isomerase and dithiol oxidase activity. The crystal structure reveals a trimeric arrangement for PmScsCΔLinker with the catalytic domains closely packed against each other. Conversely, SAXS analysis indicated that the oligomeric state of PmScsCΔLinker is concentration-dependent, with a monomeric form present at low concentrations and a dimeric form dominating at higher concentrations. For comparison, we analysed a concentration series of full-length native PmScsC by SAXS showing that the dominant species of the native protein in solution is trimeric, even at very low concentrations.

**Figure 1:**
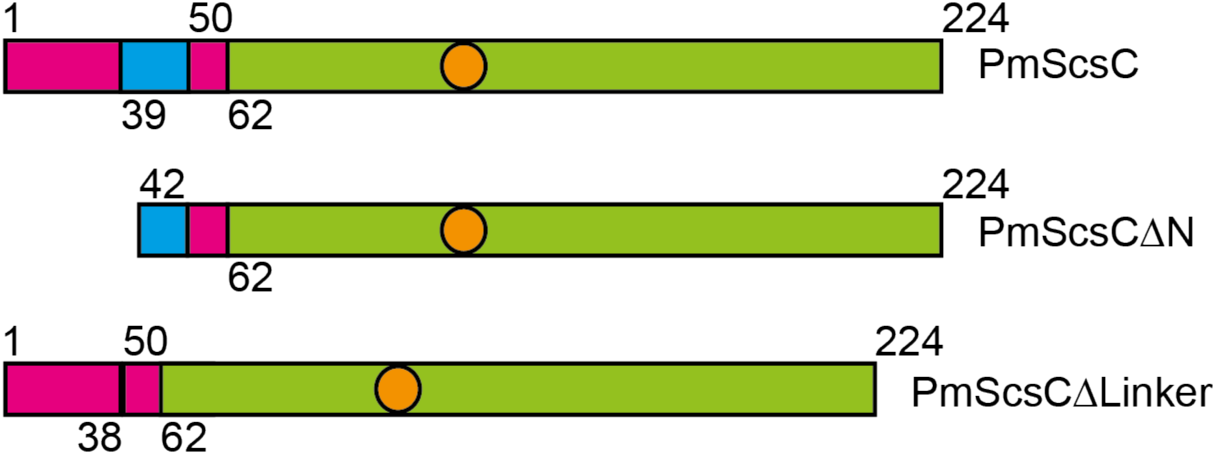
Schematic representation of the PmScsC variants used in this study. The region corresponding to the catalytic domain is shown in green, the trimerisation stem is coloured magenta and the region representing the flexible linker is shown in cyan. The orange circle represents the approximate position of the redox active cysteines.

## 2. Materials and methods

### 2.1 Molecular biology, protein expression and purification

The constructs for the expression of PmScsC and PmScsCΔN were created previously (Furlong *et al.*, 2017) using the UniProt sequence B4EV21, without the predicted periplasmic signaling sequence (i.e. residues 22-243). To make the PmScsCΔLinker expression construct, the DNA encoding the flexible peptide linker (residues 39-KADEQQAQFRQ-49) in PmScsC was deleted from the *P. mirabilis scsC* gene using overlap extension PCR, with the primers 5’-GCAATCATGGCTCTGCAGACGAAAGCACTGGCTAGCGAACATGATGCC and 5’- GGCATCATGTTCGCTAGCCAGTGCTTTCGTCTGCAGAGCCATGATTGC. Ligation independent cloning was then used to insert the mutated gene into pMCSG7. Sequencing of the construct confirmed that the encoded protein was in-frame with an N-terminal His_6_-tag and TEV protease cleavage site and revealed that residue 6 of PmScsCΔLinker was also mutated from asparagine to lysine. The protein residues are numbered based on the sequence of full-length PmScsC, after TEV protease cleavage (starting S_1_N_2_A_3_…). All proteins were expressed in *E. coli* and purified using immobilized metal affinity, TEV cleavage and size exclusion chromatography as previously described (Furlong *et al.*, 2017). SDS-PAGE with Coomassie stain was used to estimate that the purity of each protein was >95%. The concentration of the proteins was determined by measuring the A_280_ of the samples and adjusting for theoretical A_280_ of a 1 mg/ml solution of the protein (Abs 0.1%). The theoretical Abs 0.1% values determined from Protparam (Gasteiger *et al.*, 2005) were 0.528, 0.643 and 0.558 for PmScsC, PmScsCΔN and PmScsCΔLinker, respectively.

### 2.2 Disulfide isomerase and dithiol oxidase assays

The disulfide isomerase and dithiol oxidase activity of PmScsCΔLinker was determined as described previously (Furlong *et al.*, 2017). Briefly, to determine the disulfide isomerase activity, the ability of the protein to refold scrambled RNase A (scRNase A) was assessed. 10 μM PmScsCΔLinker was incubated with 40 μM scRNase A and at various time points, samples were taken and the RNAse A activity was measured in a spectrophotometric assay with a final concentration of 3 mM cytidine 3’,-5’-cyclic monophosphate (cCMP). Results are reported as a percentage of the activity of the RNase only control and are adjusted for the scrambled RNase A only negative control. The dithiol oxidase activity of PmScsCΔLinker was assessed by measuring the protein’s ability to form a disulfide bond between the two cysteines in a synthetic peptide, over time. The peptide substrate has 1,4,7,10-tetraazacyclododecane-1,4,7,10-tetraacetic acid (DOTA) bound to europium at the N-terminus and lysine methoxy-coumarin amide (MCA) at the C-terminus. As the cysteines become oxidised, the termini of the peptide are brought within close proximity, which causes an increase in fluorescence. PmScsCΔLinker (80 nM) was incubated with 8 μM of the peptide substrate and the increase in fluorescence over time was monitored using a Synergy H1 Hybrid plate reader (BioTek, Vermont, USA) reading fluorescence at excitation and emission wavelengths of 340 nm and 615 nm, respectively. The rate of peptide oxidation is reported relative to the activity of the prototypical dithiol oxidase EcDsbA. The disulfide isomerase and dithiol oxidase assays were repeated two and three times, respectively and the mean and standard deviation for each is reported.

### 2.3 Crystallisation

Initial crystallisation hits for both PmScsCΔN and PmScsCΔLinker were identified in 96-well hanging-drop vapour-diffusion experiments using commercial crystallisation screens, set up with a Mosquito robot (TTP LabTech Ltd, Melbourn, UK) at the University of Queensland Remote Operation Crystallisation and X-ray (UQ ROCX) Diffraction Facility. All crystallisation experiments were incubated at 293 K and monitored using a Rock Imager with Rock Maker software (Formulatrix Inc., Waltham, Massachusetts, USA).

Well diffracting, rod-shaped crystals of PmScsCΔN were obtained in a 96-well screening plate in a drop containing 200 nL of the PEGRx™ HT (Hampton Research, Aliso Viejo, California, USA) C10 condition, 0.1 M Sodium acetate trihydrate pH 4.5, 30% w/v PEG-mono methylether 5000, and 200 nL of 38 mg/ml PmScsCΔN in 10 mM HEPES pH7.4. Perfluoropolyether (Hampton Research, Aliso Viejo, California, USA) was used to cryoprotect the crystals before they were flash frozen in liquid nitrogen.

Needle-like crystals of PmScsCΔLinker were obtained in well F12 of the commercial screen JCSG-*plus*™ (Molecular Dimensions Limited, New-Market, UK) and then optimised in a 24-well plate grid screen around the original condition. This optimisation resulted in more robust and well diffracting crystals. Data were collected from a crystal grown in 1 μL of 0.1 M HEPES pH 7, 32% v/v Jeffamine M-600 and 1 μL of 40 mg/mL PmScsCΔLinker in 10 mM HEPES pH 7.4, 150 mM NaCl. The crystal was cryoprotected in the crystallisation condition plus 20% ethylene glycol before being flash cooled in a cryostream (100 K).

### 2.4 Data collection, structure solution and refinement

Data for the PmScsCΔN structure were collected using a wavelength of 0.95370 Å at 100 K on the Australian Synchrotron MX2 beamline using the BluIce software (McPhillips *et al.*, 2002). The data were processed and scaled into the trigonal space group P312 using XDS (Kabsch, 2010), Pointless (Evans, 2006) and SCALA (Evans, 2006). Phases were obtained by molecular replacement in Phaser (McCoy *et al.*, 2007), using *Salmonella enterica* serovar Typhimurium ScsC (PDB: 4GXZ) as the model. The initial model was improved by iterative model building in Coot (Emsley *et al.*, 2010) and refinement in PHENIX (Adams *et al.*, 2010). The quality of the final PmScsCΔN model was assessed using MolProbity (Chen *et al.*, 2010) and had an R_free_ value of 21.5% and R_work_ value of 19.6%, at a resolution of 2.15 Å. The coordinates and structure factors are deposited to the Protein Data Bank under the code 4YX8.

Data for the PmScsCΔLinker structure were collected at UQ ROCX under cryogenic conditions (100 K) using a Rigaku FR-E Superbright X-ray generator, producing X-rays at a wavelength of 1.54187 Å, and a Rigaku Saturn 944 CCD area detector. The data collection was performed with the Rigaku CrystalClear (v 2.0) program. Data were processed in iMosflm (v 7.1.3) (Battye *et al.*, 2011) with the space group H32 (also known as R32:h) and scaled in Aimless (Evans & Murshudov, 2013) as implemented in the CCP4 suite (v 6.5.008) (Winn *et al.*, 2011). Due to detector distance constraints at UQ ROCX, diffraction data were limited to a highest resolution of 2.08 Å. The crystals diffracted strongly, resulting in very high mean I/σI values (27.2 overall and 11.8 in the highest resolution shell). The structure was phased by molecular replacement using Phaser (McCoy *et al.*, 2007) and the catalytic domain of the compact PmScsC crystal structure (residues 47-224, PDB: 4XVW) was used as the search model. The trimerisation domain (without the 11 amino acid flexible linker) was then manually built into the electron density in Coot (Emsley *et al.*, 2010). The structure was refined using rounds of phenix.refine and manual adjustment in Coot, with reference to MolProbity (Chen *et al.*, 2010). The final R_free_ and R_work_ values are 20.80% and 17.55% respectively, at a resolution of 2.08 Å. The coordinates and structure factors are deposited with the Protein Data Bank under the code 6MHH. All structural figures were made in PyMOL (*The PyMOL Molecular Graphics System, Version 1.6*) and structural alignments were performed in Coot using the superpose command (Emsley *et al.*, 2010) which aligns Cαs by least squares. The Contact program from the CCP4 suite (Winn *et al.*, 2011) was used to analyse the interactions between the trimerisation stems of PmScsC/PmScsCΔLinker.

### 2.5 SAXS

SAXS data were collected on the SAXS-WAXS beamline at the Australian Synchrotron (Kirby *et al.*, 2013). Data reduction was carried out using Scatterbrain software (v 2.71) (*Scatterbrain - Software for acquiring, processing and viewing SAXS/WAXS data at the Australian Synchrotron*), and corrected for solvent scattering, sample transmission, and detector sensitivity. The 0.1, 0.5, 2.0, 5.0 and 10.0 mg/mL samples of PmScsCΔN were made by diluting a 30 mg/mL stock in 10 mM HEPES pH 7.4, 150 mM NaCl. For PmScsC and PmScsCΔLinker, samples were taken throughout centrifugal concentration and the concentration of these samples was adjusted to 0.1, 0.5, 2.0, 5.0 or 10.0 mg/mL in 10 mM HEPES pH 7.4, 150 mM NaCl. These samples, along with the blank (10 mM HEPES pH 7.4, 150 mM NaCl) were loaded into a 96-well plate (Table 2) (Trewhella *et al.*, 2017). The estimated molecular mass was calculated from the Porod volume (Fischer *et al.*, 2010) and from *I*(0) using contrast and partial specific volumes determined from the protein sequences using MULCh (v 1.1) (Whitten *et al.*, 2008). For PmScsCΔN, data processing and Guinier analysis was performed using Primus (v 3.2) (Konarev *et al.*, 2003). The pair-distance distribution function (*p*(*r*)) was generated from the experimental data using *GNOM* (v 4.6) (Svergun, 1992), from which *I*(0), *R*_g_ and *D*_max_ were determined. The program *DAMMIN* (v 5.3) (Svergun, 1999) was used to generate 16 dummy-atom models for the protein, which were averaged using the program *DAMAVER* (v 2.8.0) (Volkov & Svergun, 2003), and the resolution of the averaged structures was estimated using SASRES (Tuukkanen *et al.*, 2016). All 16 dummy-atom models were used in the averaging procedure. The model scattering curve for PmScsCΔN (SASBDB ID: SASDEK4) was calculated from the crystal structure (PDB ID: 4YX8) using CRYSOL (v 2.8.3) (Svergun *et al.*, 1995).

For PmScsC (SASBDB ID: SASDER4) and PmScsCΔLinker (SASBDB ID: SASDEQ4), data were modelled as a linear combination of model scattering curves to quantify the proportion of different oligomeric states in solution (Hu *et al.*, 2012). The structure of PmScsC monomer (chain ID: A), dimer (chain ID: A, B) and trimer (chain ID: A, B, C) were taken from a CORAL model previously optimized against SAXS data for PmScsC (SASDB94) (Furlong *et al.*, 2017). The structure of PmScsCΔLinker monomer (chain ID: A), dimer (chain ID: A, B) and trimer (chain ID: A, B, C) were taken from the crystal structure of PmScsCΔLinker (PDB ID: 6MHH). Scattering curves for each oligomer were calculated using CRYSOL (v 2.8.3) (Svergun *et al.*, 1995). The calculated scattering curves (with units of *e*^2^) were multiplied by *r*_e_^2^(cm^2^ *e*^-2^) × *N*_A_ (mol^-1^) × 10^-3^ (L cm^-3^) = 4.78181 × 10^-5^ (cm^-1^ *e*^-2^ mol^-1^ L). This normalization permits the coefficients arising from a fit to experimental data on an absolute scale to be interpreted as the molar concentration of each oligomer present in solution. For PmScsC, a linear combination of monomer and trimer scattering curves were fit to the scattering data at each concentration. To reduce the number of free parameters, the concentration of monomer for each SAXS curve was optimized together with a common equilibrium constant, 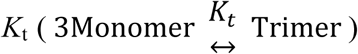, and the concentration of trimer was calculated from *K*_t_ and the momomer concentration. For PmScsCΔLinker, a linear combination of monomer, dimer and trimer scattering curves were fit to the scattering data at each concentration. To reduce the number of free parameters, the concentration of monomer for each SAXS curve was optimized together with two common equilibrium constants, 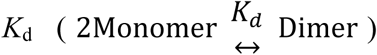, and 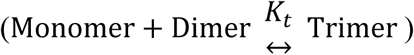. The concentration of dimer was calculated from *K*_d_ and the monomer concentration, while the concentration of trimer was calculated from *K*_t_ and the concentration of monomer and dimer.

## 3. Results

### 3.1 PmScsCΔN crystal structure and SAXS analysis

The design and biochemical characterisation of PmScsCΔN has been reported previously (Furlong *et al.*, 2017). The present work reports the crystal structure of this variant. Crystals of the PmScsCΔN mutant diffracted to a resolution of 2.15 Å at the Australian Synchrotron. The crystal structure of the mutant was solved using molecular replacement and there was only one PmScsCΔN molecule in the asymmetric unit (Table 1, Figure 2A, B). In the crystal packing, PmScsCΔN makes contact with 6 symmetry related molecules, but none involve packing against the catalytic motif or the positively charged surface patch surrounding it. The structure of PmScsCΔN is very similar to the structure of the catalytic domain present in each of the three previously reported PmScsC structures (Figure 2C, RMSD: 0.51-0.68 Å, Cα atoms of 174 residues aligned, between chain A of each structure). Small-angle scattering (Table 2) was used to determine the low-resolution solution structure of PmScsCΔN (Figure 3). The scattering data for PmScsCΔN is representative of a monomeric globular particle and is consistent with the scattering curve calculated from the crystal structure of PmScsCΔN (PDB ID: 4YX8). The crystal structure also shows good agreement with the dummy atom model derived from the scattering data (Figure 3C).

**Figure 2:**
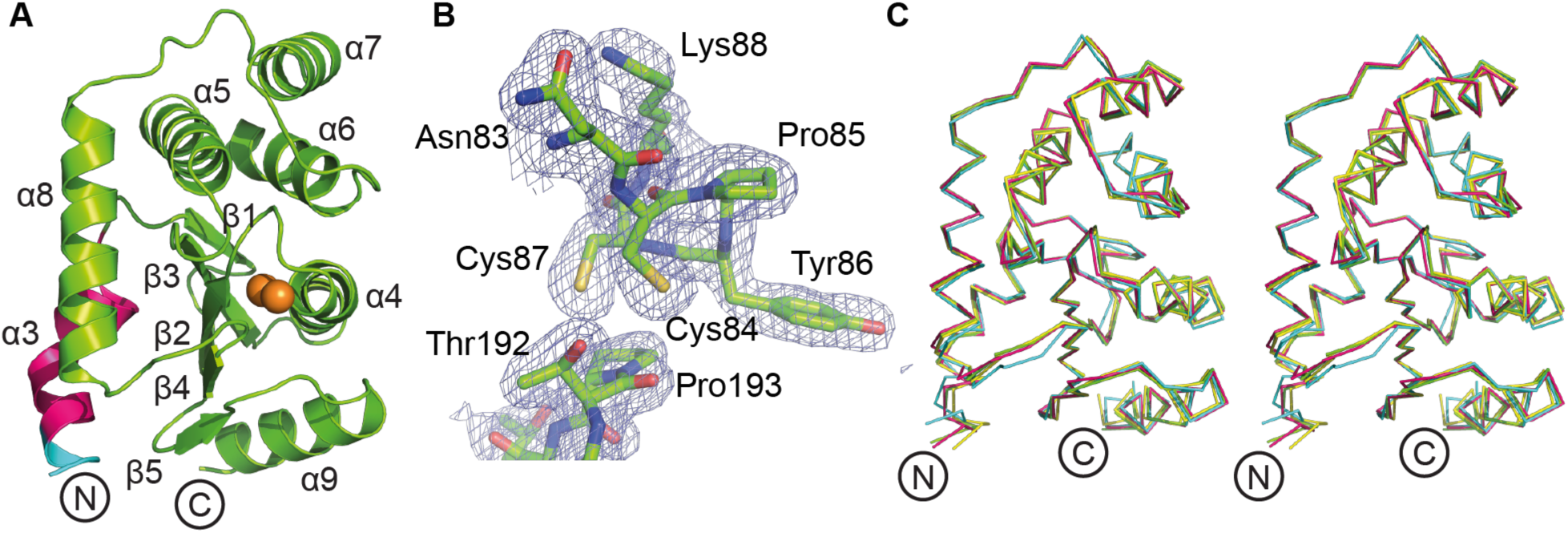
Structural characterisation of PmScsCΔN. **A.** Structure of the PmScsCΔN variant. Structural features are coloured as in Figure 1 and the secondary structure elements are labelled based on the native PmScsC structures (Furlong *et al.*, 2017). **B.** Electron density of the catalytic region in PmScsCΔN. The wire mesh represents the 2F_o_-F_c_ electron density map (generated in phenix.maps) contoured to 1σ and shown within a 2 Å radius of each atom. **C.** The structural alignment of PmScsCΔN (green) with residues 46-221 of chain A from the compact (cyan, PDB ID: 4XVW), transitional (yellow, PDB ID: 5IDR) and extended (magenta, PDB ID: 5ID4) PmScsC structures (Furlong *et al.*, 2017) (RMSD: 0.51-0.68 Å, 174 residues aligned). This superimposition shows that the structures are similar whether or not the trimerisation domain is present.

**Figure 3:**
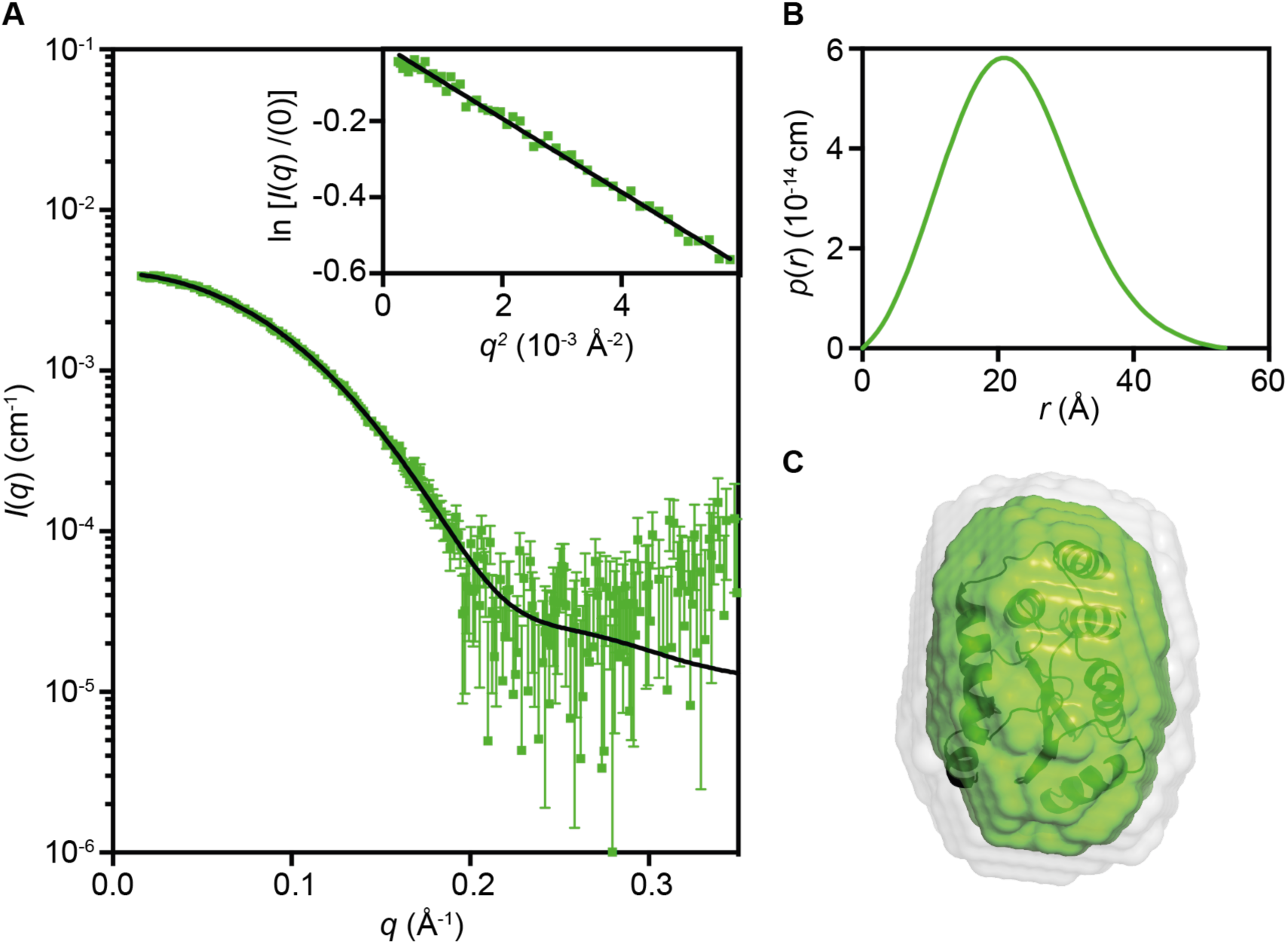
Small angle X-ray scattering data for PmScsCΔN. **A**. Measured scattering data for PmScsCΔN. The scattering profile of rigid-body models are shown as solid black lines overlaid on the scattering data for PmScsCΔN (*χ*^2^ = 1.12; CorMap test (Franke *et al.*, 2015), 302 points, *C*=14, *P*=0.018). Inset: Guinier plot for PmScsCΔN (*R*^2^ = 0.991) **B.** The pair-distance distribution function, *p*(*r*), derived from the scattering data are indicative of a globular structure for PmScsCΔN (multiplied by a factor of 4 for clarity) with a maximum dimension of ~55 Å. **C.** Probable shape of PmScsCΔN obtained from the filtered average of 16 dummy-atom models (green envelope): *χ*^2^ = 1.079 ± 0.001; NSD = 0.522 ± 0.007; Resolution = 19 ± 2. Images in C were generated using PyMol, where the grey shapes represent the total volume encompassed by the aligned dummy-atom models and the corresponding rigid-body model is shown aligned to the filtered model.

**Table 1:**
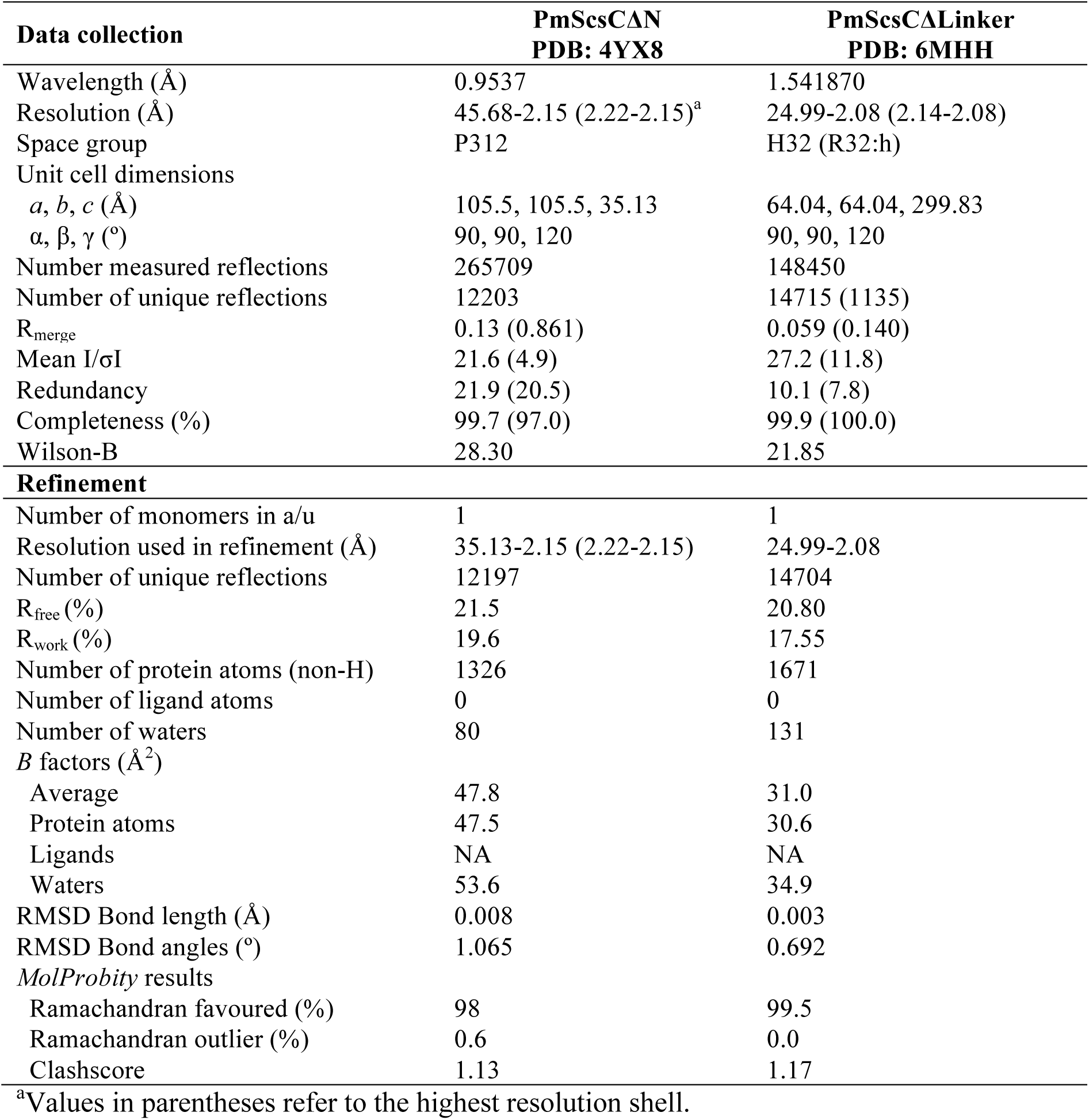
Crystallography statistics for PmScsCΔN and PmScsCΔLinker.

### 3.2 Activity of PmScsCΔLinker

A key structural feature of native PmScsC is an 11 amino acid flexible linker, situated in the trimerisation stem and linking to the catalytic domain. This short region facilitates the conformational variations observed in the crystal structures and the dynamic motion of the trimeric protein observed in SAXS analyses (Furlong *et al.*, 2017). To better understand the role of the flexible linker we designed a PmScsCΔLinker variant in which the 11-residue linker is deleted, and characterised its function in standard assays. We first assessed the activity of PmScsCΔLinker in the scrambled RNase A assay and found that it was poorly active, with baseline activity like that of a typical dithiol oxidase such as EcDsbA and PmScsCΔN (Figure 4A). We then assessed its activity in the model peptide dithiol oxidase assay, in which EcDsbA and PmScsCΔN are both active. However, unlike the monomeric proteins EcDsbA and PmScsCΔN, PmScsCΔLinker was also inactive in the dithiol oxidase assay (Figure 4B). Disulfide isomerase activity is thought to be associated with oligomerisation of thioredoxin-fold proteins (Furlong *et al.*, 2017, Ke *et al.*, 2006, Sun & Wang, 2000) so we were interested to determine whether PmScsCΔLinker forms a trimer - like the full-length native protein - or a monomer - like the truncated PmScsCΔN variant.

**Figure 4:**
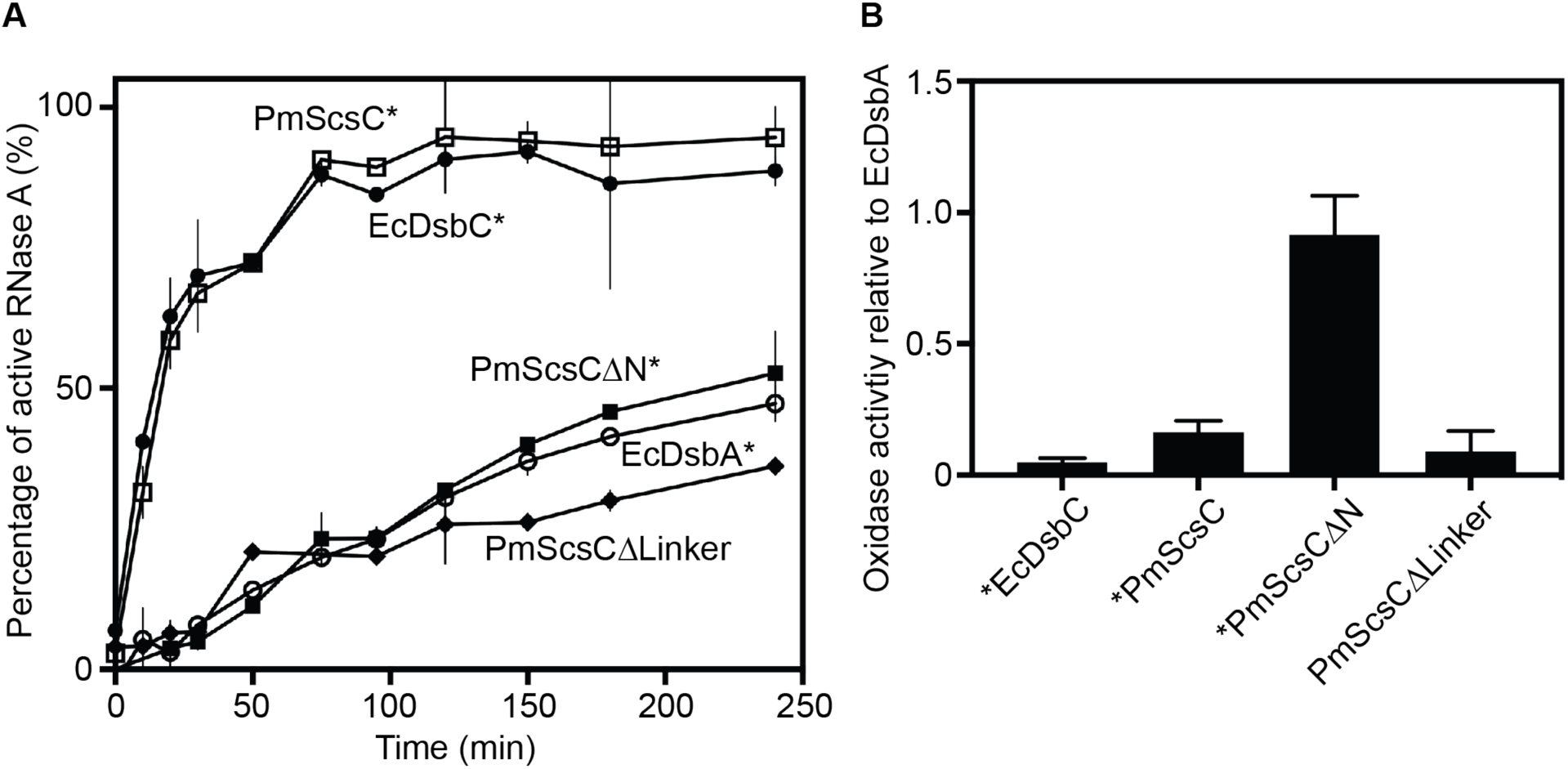
PmScsCΔLinker biochemical activity. **A.** The ability of PmScsCΔLinker to isomerise the incorrect disulfide bonds in scrambled RNase A is poor – i.e. even lower than that of the dithiol oxidase EcDsbA. **B.** PmScsCΔLinker has poor dithiol oxidase activity; i.e. much less than monomeric PmScsCΔN and similar to that of protein disulfide isomerases EcDsbC and PmScsC. Asterisks indicate data published previously (Furlong *et al.*, 2017). Error bars on both panels represent the standard deviation of the mean.

### 3.3 The crystal structure reveals a trimeric form of PmScsCΔLinker

The three crystal structures of native PmScsC reported previously show that the linker residues can adopt helical (extended conformation), strand (intermediate conformation) or loop (compact conformation) secondary structure (Furlong *et al.*, 2017). In the extended conformation crystal structure, the linker adopts a helical conformation thereby creating a very long helix (35 residues, approximately 9.5 turns, Figure 5A) extending from the trimerisation stalk, through into the catalytic domain of PmScsC.

**Figure 5:**
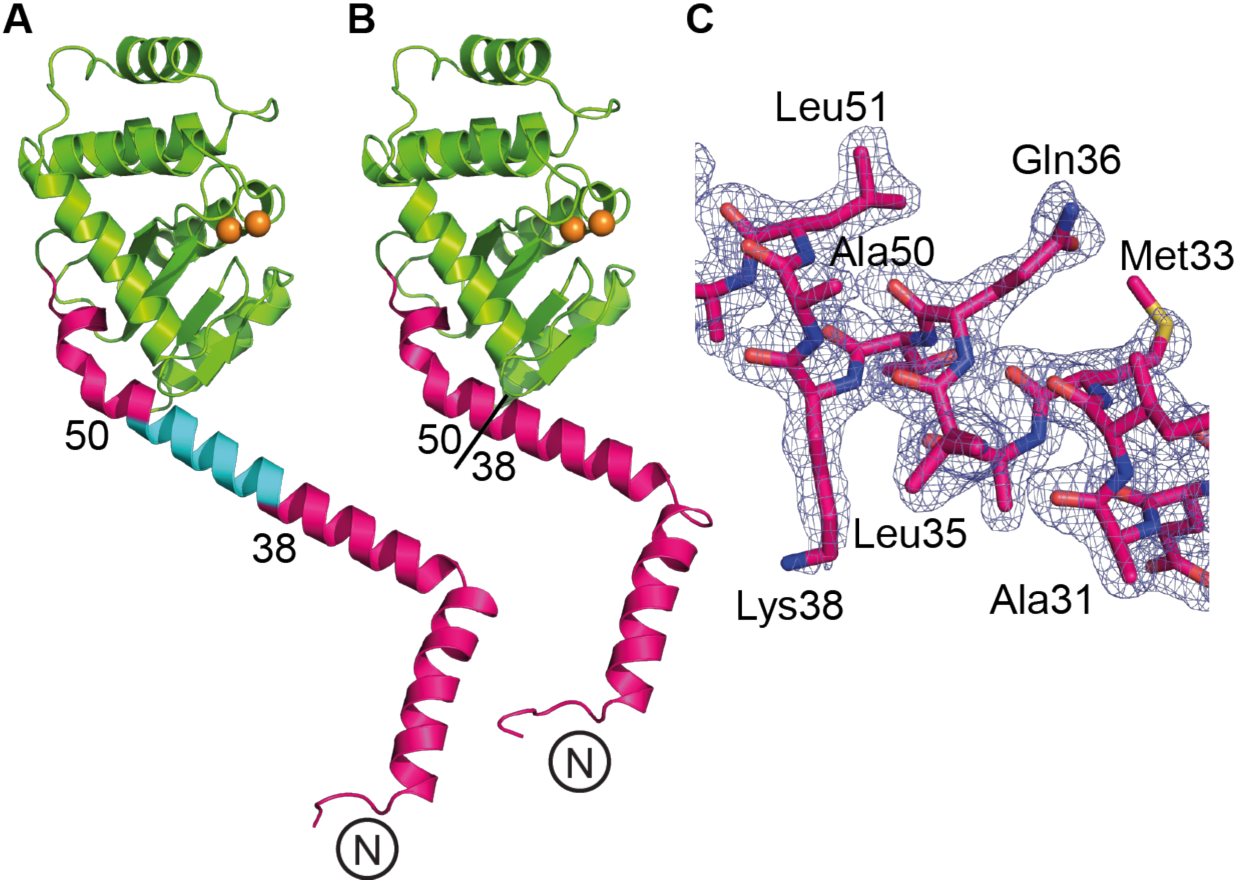
PmScsCΔLinker crystal structure. **A.** Single protomer of the extended PmScsC (PDB: 5ID4). **B.** Single protomer of PmScsCΔLinker. The different features of the crystal structures are coloured as in Figure 1. The N-terminus of each structure is labelled and the positions of the residues flanking the flexible linker in the native structure are indicated. **C.** Electron density around the deleted region in the PmScsCΔLinker structure. The wire mesh represents the 2F_o_-F_c_ electron density map (generated in phenix.maps) contoured to 1σ and shown within a 2Å radius of each atom.

We solved the crystal structure of PmScsCΔLinker at a resolution of 2.08 Å using molecular replacement (Table 1, Figure 5B, C). The variant crystallised in the same space group (H32) and crystallisation condition as the extended conformation PmScsC (PDB: 5ID4). However, the unit cell dimensions of PmScsCΔLinker are shorter than those of the extended PmScsC structure (64.0 Å, 64.0 Å, 299.8 Å compared to 86.7 Å, 86.7 Å, 330.0 Å). The crystal structure of PmScsCΔLinker– like the crystal structure of the extended form of PmScsC – is trimeric through crystallographic symmetry, with a single protomer in the asymmetric unit (Figure 6A). The regions on either side of the linker region reported to be helical in the three native PmScsC crystal structures are directly connected in PmScsCΔLinker and are also helical.

**Figure 6:**
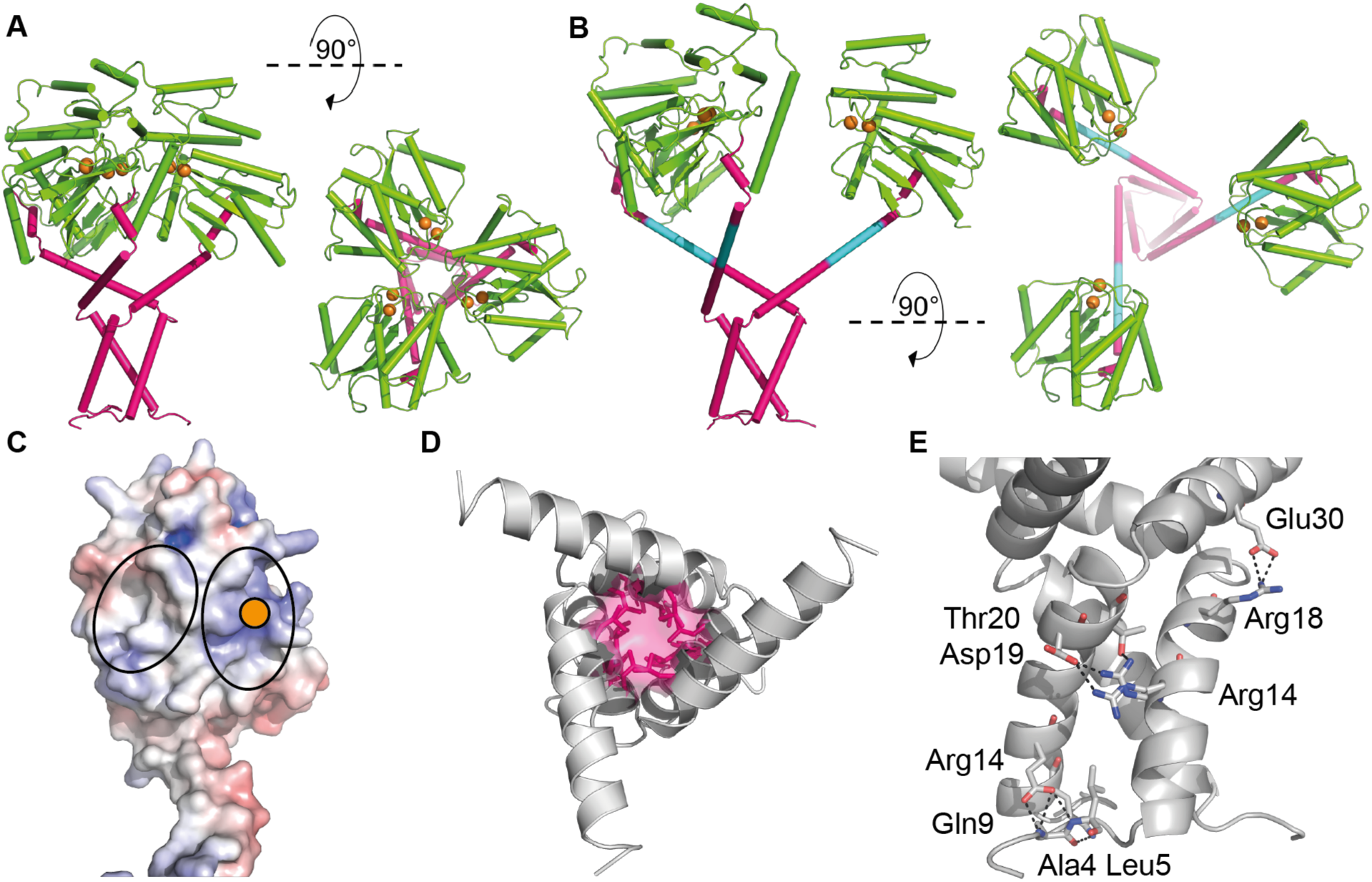
Trimerisation of PmScsCΔLinker. **A.** Side and top views of the catalytic domain of the crystal structure of PmScsCΔLinker. The crystal structure reveals a trimer through crystallographic symmetry. In many respects this crystal structure resembles that of native extended PmScsC **B** (PDB: 5ID4). However, the extended helix linking the trimerisation domain to the catalytic domain is shorter in PmScsCΔLinker due to deletion of the linker (cyan in **B**). In both **A** and **B**, the proteins are coloured using the scheme from Figure 1. **C.** The electrostatic surface potential of the catalytic domain of the PmScsCΔLinker protomer structure. Electrostatic calculations were performed using APBS (Jurrus *et al.*, 2018) and contoured to -7.5 (red) and +7.5 kT/e (blue). A solid orange circle indicates the position of the catalytic cysteines and black ovals surround the regions of the catalytic domains in close contact with adjacent catalytic domains in the PmScsCΔLinker crystal structure. **D.** The hydrophobic core of the trimerisation stem of PmScsCΔLinker; sidechains of residues contributing to the hydrophobic core are coloured pink **E.** Amino acid residues mediating side chain polar and electrostatic interactions between the trimerisation stems of two PmScsCΔLinker protomers. Black dashed lines represent the hydrogen bonds and those residues involved are labelled.

The crystal structure of PmScsCΔLinker is similar to that of extended PmScsC except for a shortening of the long helix by almost three turns (Figure 5A, B). This shortened helix in the crystal trimer of PmScsCΔLinker brings the three catalytic domains into much closer proximity in the variant compared with the native extended conformation. This proximity is evident from the distance between the catalytic cysteines of protomers: 15 Å in PmScsCΔLinker and 34 Å in the extended native structure (measured between the Cβ atoms of C84, Figure 6A, B).

The PmScsCΔLinker crystal structure also reveals interactions between each of the catalytic domains of the trimer. The surface interface is formed between a positively charged patch close to the CXXC motif on the catalytic domain of one protomer and a neutral/hydrophobic patch on another (Figure 6C). This interface is not formed in the extended PmScsC structure, which has both of these regions exposed to solvent and it is also not formed by crystal packing in the PmScsCΔN structure.

The trimerisation stems of all three native PmScsC crystal structures are α-helical with a hydrophobic core, and hydrogen bonds between surface exposed polar/charged residues. These features are conserved in the crystal structure of PmScsCΔLinker (Figure 6D and E). However, it should be noted that PmScsCΔLinker has an additional unanticipated mutation (Asn6Lys) at the N-terminus. In the three native crystal structures the side chain of the Asn6 residue forms an intramolecular hydrogen bond to the main chain of Gln9. In the PmScsCΔLinker crystal structure, the side chain of Gln9 adopts the same conformation as in all three native PmScsC structures but the mutated residue Lys6 does not form an equivalent hydrogen bond with the Gln9 main chain. The lack of this specific hydrogen bond could potentially increase N-terminal disorder, though the crystal structure suggests there is little impact on the hydrogen bonds and hydrophobic contacts that contribute to trimerisation.

### 3.4 SAXS reveals that the oligomeric state of PmScsCΔLinker is concentration dependent

We had previously observed that the PmScsC RHP variant (in which the flexible linker is replaced with a rigid helix) forms a trimer - like the native protein - but is inactive in the protein disulfide isomerase assay (Furlong *et al.*, 2017). This lack of activity was thought to be a consequence of reduced catalytic domain motion observed in solution using small angle scattering (Furlong *et al.*, 2017). This finding led to the hypothesis that potent protein disulfide isomerase activity requires at least two catalytic domains (e.g. dimer, trimer) which have the ability to move independently of each other (Furlong *et al.*, 2017). Our present results show that PmScsCΔLinker also forms a trimer and is also inactive as a protein disulfide isomerase. We therefore anticipated that the lack of protein disulfide isomerase activity was a consequence of reduced catalytic domain rigid body motion, and sought to confirm this in solution using small angle scattering.

Unexpectedly, initial SAXS data for PmScsCΔLinker displayed a strong dependence on concentration, consistent with an equilibrium between different PmScsCΔLinker oligomeric forms (monomer, dimer, trimer). This result raised the possibility that, at low concentrations, the wild-type protein may also be present as a mixture of monomer, dimer and trimer, and this might also impact on its function or its regulation. To investigate this possibility further, SAXS data were collected from native PmScsC (Figure 7) and PmScsCΔLinker (Figure 8) at concentrations between 0.1 and 10.0 mg/mL (Table 2). At the lowest concentration (0.1 mg/mL), the shape of the *p*(*r*) function for native PmScsC differed from the shapes of those at higher concentrations, indicating some dissociation of the wild-type protein. This effect was quantified by fitting a linear combination of monomer, dimer and trimer scattering to the data at each concentration (Figure 7 and Table 3). For PmScsCΔLinker, the results are very different; the shape of the *p*(*r*) curve changes significantly with each concentration evaluated. Again, this effect was quantified by fitting a linear combination of monomer, dimer and trimer scattering to the data at each concentration (Figure 8 and Table 3). The relative amounts of each were constrained by common equilibrium constants. The analysis worked well, but was limited by the fact that the structures of monomer and dimer forms in solution are not known. In each case, we made a simplistic assumption that a monomer adopted the structure of a single protomer from the corresponding crystal structure, and a dimer could be represented by removing a single protomer from the trimer crystal structure. We also know that native PmScsC is dynamic in solution (Furlong *et al.*, 2017). While the fits we obtained of the model curves to the scattering data are reasonable, there are systematic differences between the experimental data and the model, especially at higher concentrations. The limitations of the modeling (lack of accurate models, dynamic motion) explains most of this variation.

**Figure 7:**
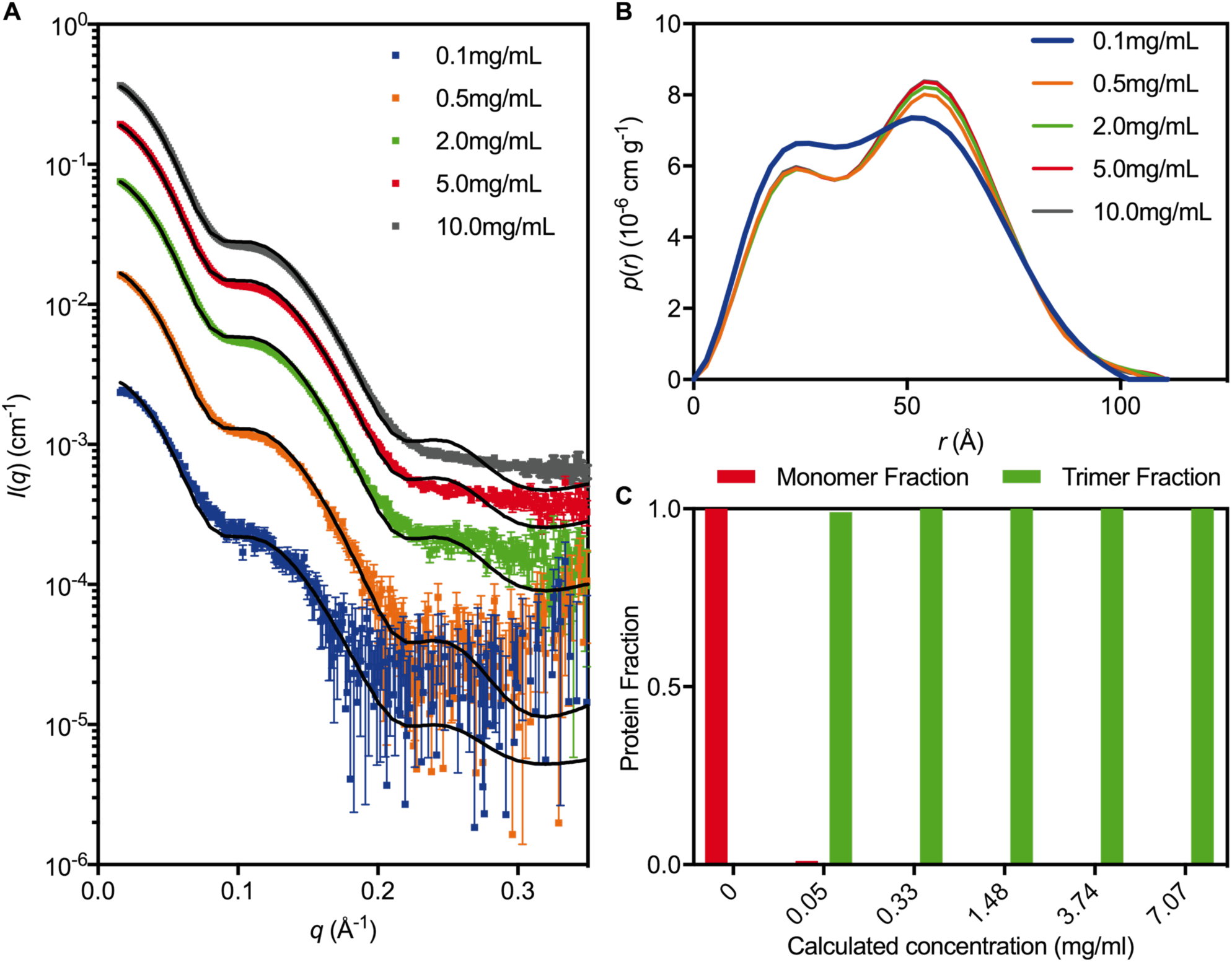
Native PmScsC SAXS data. **A.** Scattering data collected at ~0.1 mg/mL (blue, *χ*^2^ = 2.55), ~0.5 mg/mL (orange, *χ*^2^ = 7.05), ~2.0 mg/mL (green, *χ*^2^ = 72.85), ~5.0 mg/mL (red, *χ*^2^ = 371.94), ~10.0 mg/mL (grey, *χ*^2^ = 815.23). The overlaid line (black) on each scattering curve is the fitted linear combination of monomer and trimer scattering curves. **B.** The pair-distance distribution function, *p*(*r*), derived from the scattering data normalised by concentration show the 0.1 mg/mL data set differs slightly from that of the other concentrations (which all overlay), indicating the possibility that monomer is present in solution at the lowest concentration. **C.** The fraction of protein present as monomer and trimer, derived from the fitted models, shows that at total protein concentrations above 10 μM, the protein is almost exclusively present as a trimer in solution.

**Figure 8:**
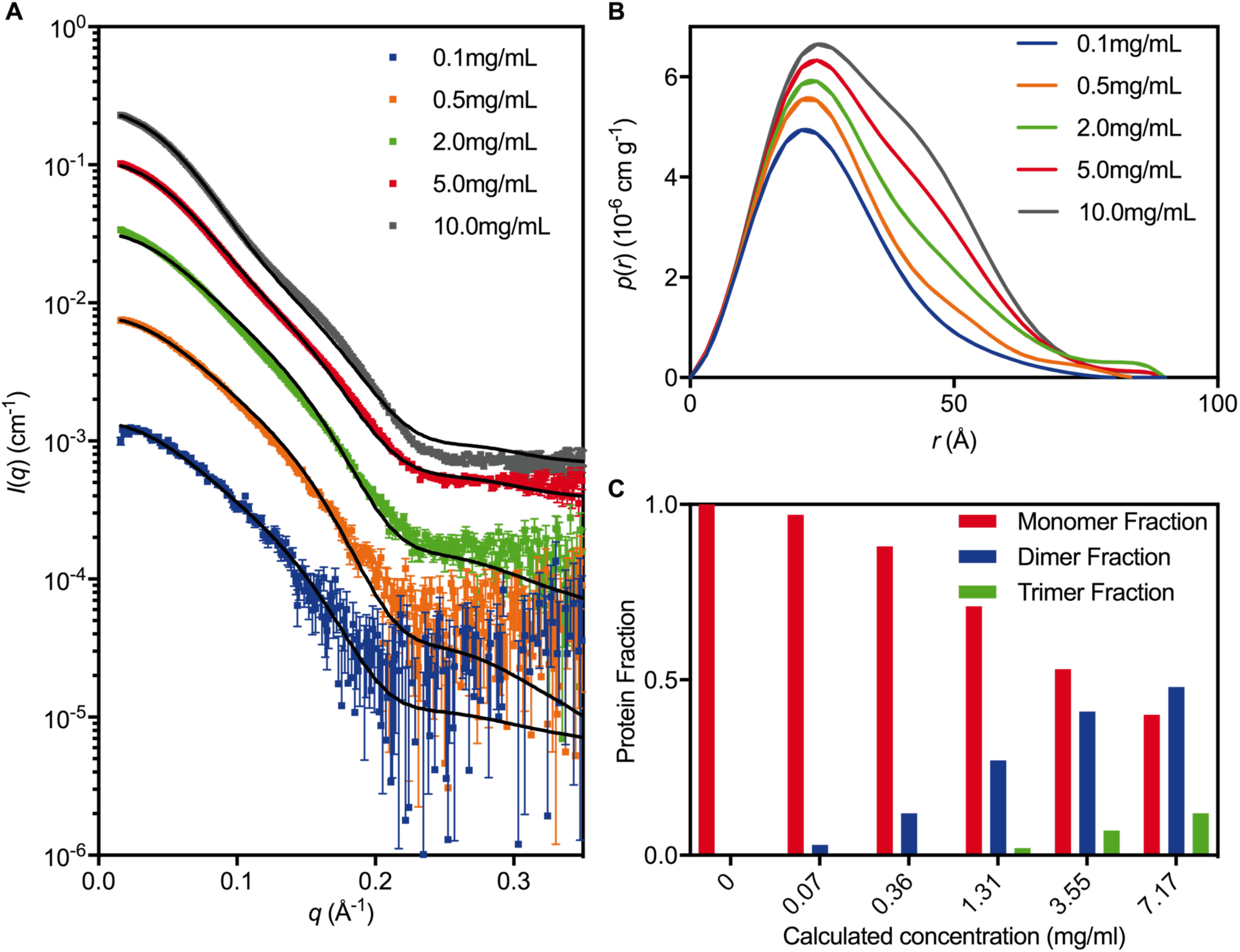
PmScsCΔLinker SAXS data. **A.** Scattering data collected at ~0.1 mg/mL (blue, *χ*^2^ = 1.27), ~0.5 mg/mL (orange, *χ*^2^ = 2.19), ~2.0 mg/mL (green, *χ*^2^ = 6.89), ~5.0 mg/mL (red, *χ*^2^ = 37.59), ~10.0 mg/mL (grey, *χ*^2^ = 177.29). The overlaid line (black) on each scattering curve is the fitted linear combination of monomer and trimer scattering curves. **B.** The pair-distance distribution function, *p*(*r*), derived from the scattering data normalised by concentration show that the oligomeric composition of the solution is highly dependent on concentration. **C.** The fraction of protein present as monomer, dimer, and trimer, derived from the fitted models, shows the monomeric species is dominant in solution until the total protein concentration reaches ~240 μM. The concentration of trimer rises slowly, and reaches a protein fraction of ~0.2 at 1.7 mM (40 mg/mL), which is the concentration at which the crystallisation was performed.

**Table 2:**
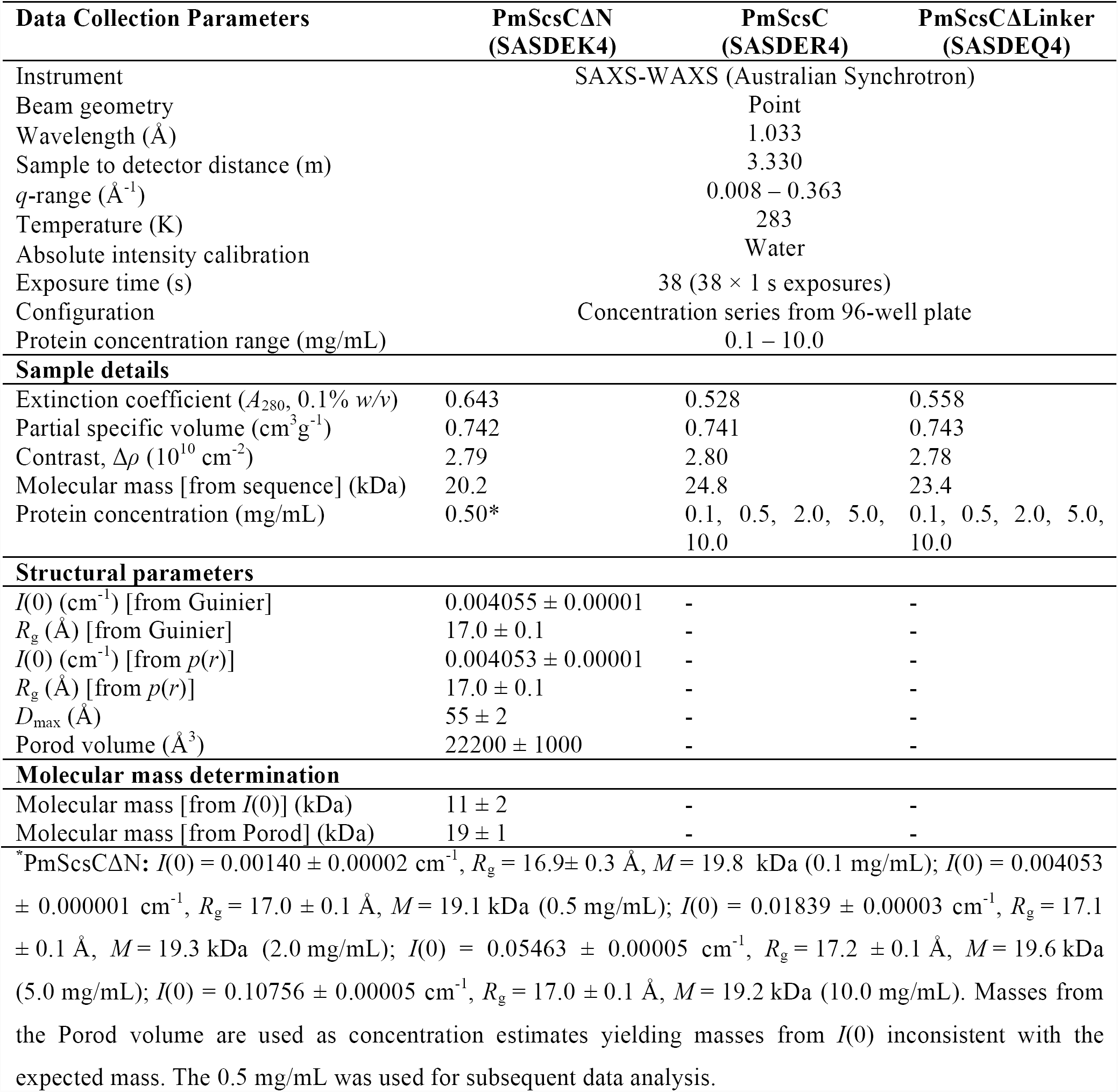
SAXS data collection and analysis details.

**Table 3:** Oligomer fractions and equilibrium constants

| Measured<br>Concentration<br>(mg/mL) | Calculated<br>Concentration<br>(mg/mL) | Monomer<br>Fraction | Dimer<br>Fraction | Trimer<br>Fraction |
| --- | --- | --- | --- | --- |
| *Native PmScsC* ($K_t = 6.3 \times 10^{-18} \text{ M}^2$) | | | | |
| 0.1 | 0.05 | 0.01 | - | 0.99 |
| 0.5 | 0.33 | 0.00 | - | 1.00 |
| 2.0 | 1.48 | 0.00 | - | 1.00 |
| 5.0 | 3.74 | 0.00 | - | 1.00 |
| 10.0 | 7.07 | 0.00 | - | 1.00 |
| *PmScsCΔLinker* ($K_d = 2.1 \times 10^{-4} \text{ M}$; $K_t = 7.30 \times 10^{-4} \text{ M}$) | | | | |
| 0.1 | 0.07 | 0.97 | 0.03 | 0.00 |
| 0.5 | 0.36 | 0.88 | 0.12 | 0.00 |
| 2.0 | 1.31 | 0.71 | 0.27 | 0.02 |
| 5.0 | 3.55 | 0.53 | 0.41 | 0.07 |
| 10.0 | 7.17 | 0.40 | 0.48 | 0.12 |

Overall, the SAXS analysis suggests that only 1% of native PmScsC is present as monomer in solution even at the lowest concentration of 0.1 mg/mL; we conclude that the trimer is the dominant form of the native protein in solution even at low concentrations. For the variant PmScsCΔLinker, it is a different story; the protein is in equilibrium between different oligomerisation states. At 0.1 mg/mL approximately 97% of the PmScsCΔLinker solution is monomer, 3% is dimer with negligible trimer. At 10 mg/ml these proportions change to 40% monomer, 48% dimer and 12% trimer.

## 4. Discussion

This study focused on characterising two variants of PmScsC, which - in its native form - is a highly dynamic, trimeric disulfide isomerase. The native PmScsC protomer consists of an N-terminal trimerisation stem with an 11 amino acid flexible linker connected to a C-terminal thioredoxin fold catalytic domain harbouring a CXXC active site. To better understand the role of the trimerisation stem and flexible linker we designed the variants PmScsCΔN and PmScsCΔLinker. PmScsCΔN lacks the first 41 amino acids of the native PmScsC construct; its loss of disulfide isomerase activity and gain of dithiol oxidase activity has been reported previously (Furlong *et al.*, 2017). PmScsCΔLinker is newly reported here, and lacks the 11 amino acids that form the flexible linker. We reported here the structure of PmScsCΔN and analysed the structure and biochemical function of PmScsCΔLinker.

The X-ray crystal structure of PmScsCΔN agrees with previous MALLS data showing that this variant is monomeric (Furlong *et al.*, 2017). The crystal structure is also consistent with solution SAXS data and the low-resolution model obtained from SAXS analysis. Although the PmScsCΔN crystal structure is very similar to the structures of the catalytic domains in each of the three native PmScsC crystal structures, unlike the native enzyme it lacks disulfide isomerase (disulfide shuffling) activity but instead has dithiol oxidase (disulfide forming) activity. The PmScsC catalytic domain has an acidic catalytic cysteine and an energetically unfavourable disulfide (Furlong *et al.*, 2017) – that are also features of the archetypal but monomeric dithiol oxidase EcDsbA – so it is perhaps not surprising that removal of the N-terminal trimerisation residues leads to oxidase activity in a model assay. Moreover, the structural similarity between monomeric PmScsCΔN and the catalytic domains of native trimeric PmScsC supports the notion that it is the N-terminal stem – defining whether the catalytic domain is monomeric or trimeric - that is the determinant of isomerase or oxidase activity. Put simply, the differential activity we observe between native PmScsC and variant PmScsCΔN is not a specific feature of the catalytic domain because the same catalytic domain has the propensity for oxidase (monomeric protein) or isomerase (trimeric protein) activity depending on its context. The isomerase activity of PmScsC requires both its N-terminal trimerisation domain and its C-terminal catalytic domain, in the same way that the isomerase activity of EcDsbC requires its dimerisation domain and its catalytic domain (Sun & Wang, 2000).

Our SAXS analysis showed that native PmScsC is a trimer in solution, even at low concentrations. However, SAXS analysis of PmScsCΔLinker showed that it exists predominately as a monomer or a low affinity dimer (*K*_d_ ~ 200 μM) in preference to a trimer in solution even though it crystallizes in the trimeric form. Together these data suggest that removal of the flexible linker of PmScsC between the trimerisation stem and catalytic domains has the unintended consequence of reducing the propensity to form a stable trimer in solution.

The crystal structure of PmScsCΔLinker provides insight into why the trimeric form of the variant may be energetically unfavourable compared with the monomeric and dimeric forms in solution. The trimerisation stem interactions of PmScsCΔLinker are similar to those observed in the native enzyme crystal structures, but all three catalytic motifs are in closer proximity to each other than in any of the native PmScsC crystal structures. This close packing suggests the possibility of unfavorable contacts between catalytic domains in PmScsCΔLinker that may contribute to the instability of the trimer. Indeed, the PmScsCΔLinker crystal structure reveals a close association of a positively charged patch near the catalytic motif on the accessible surface of one protomer with a neutral/hydrophobic patch on another. In the crystal packing of the PmScsCΔN structure (which does not have the constraint of the trimerisation stem), there are no contacts involving the positively charged surface patch near the catalytic motif, suggesting that packing against this site is not favoured. Thus, the formation of the PmScsCΔLinker trimer is likely to be dependent on the balance of favourable interactions formed between the trimerisation stems and unfavourable interactions between the catalytic domains. At lower concentrations the unfavourable interactions between the catalytic domains may override the favourable interactions but at higher concentrations, with more molecules in close contact, it may be more thermodynamically favourable for the hydrophobic portion of the trimerisation helices to interact with each other. This interplay between interactions of the catalytic domains and trimerisation stems could explain why the oligomeric state of PmScsCΔLinker in solution alters with concentration of the protein. Taken together, the crystal structure and SAXS analysis of PmScsCΔLinker suggests that the 11 amino acid linker plays a dual role by (i) acting as a spacer between the trimerisation and catalytic domains to enable formation of a stable trimer and (ii) to provide the necessary functional dynamics and cooperativity between the catalytic domains (Furlong *et al.*, 2017).

Removal of the 11 amino acid flexible linker from PmScsC impacts on the activity of the protein: unlike the native protein, the PmScsCΔLinker has no disulfide isomerase activity. Structural characterisation of PmScsCΔLinker revealed two potential reasons for this result. Firstly, at the concentration used in the assay (10 μM) it is likely that most of the protein would be monomeric (Figure 8C), and we know from PmScsCΔN variant studies that monomeric protein has low protein disulfide isomerase activity (Furlong *et al.*, 2017). Secondly, even if there was sufficient trimeric PmScsCΔLinker present in solution, the crystal structure suggests there would be limited space between the catalytic domains to bind a misfolded protein substrate, and possibly limited motion to allow disulfides to be shuffled. It is interesting to note that PmScsCΔLinker is inactive in the dithiol oxidase assay. Given that the protein would be predominantly monomeric at the concentration tested in this assay (80 nM, Figure 8C) we expected that, like monomeric PmScsCΔN, it would have oxidase activity. One explanation for this result is that the N-terminal hydrophobic regions of PmScsCΔLinker somehow occlude the substrate-binding site that is available in monomeric PmScsCΔN.

Overall, our structural and functional characterisation of PmScsCΔN and PmScsCΔLinker provides new insight into the importance of unique structural features of the trimeric disulfide isomerase PmScsC. We confirmed previous assumptions that disulfide isomerases are similar to but differentiated from dithiol oxidases by requiring both an oligomerisation domain and a catalytic domain. We further propose that the flexible linker of PmScsC is important not just as a shape-shifting peptide, but also as a spacer, to enable functional flexibility between the catalytic domains. We suggest that this functional flexibility between the catalytic domains is necessary for cooperative enzymatic activity of disulfide isomerase PmScsC. Finally, on a more general note, we found that deleting amino acids from a protein can have unintended consequences. In this case, deletion of the flexible linker unexpectedly changed the oligomerisation kinetics of the protein, even though the region directly involved in trimerisation remained intact.

## Acknowledgments

We acknowledge the use of the University of Queensland Remote Operation Crystallisation and X-ray (UQ ROCX) facility and the support from staff, Gordon King and Karl Byriel. We also acknowledge use of the Australian Synchrotron MX and SAXS facilities and thank the staff for their support. This work was supported by an Australian Research Council Laureate Fellowship (FL0992138, to J.L.M.), an Australian Government Research Training Program stipend (to E.J.F.) and an Institute for Molecular Bioscience Research Advancement Award (to E.J.F.).

